# Targeted Reduction of Pathogenic Heteroplasmy Through Binding of G-Quadruplex DNA

**DOI:** 10.1101/358101

**Authors:** Mansur M. Naeem, Rathena Maheshan, Sheila R. Costford, Brett A. Kaufman, Neal Sondheimer

## Abstract

Pathogenic mitochondrial DNA (mtDNA) variants are typically heteroplasmic, with coexistence of variant and wild type genomes. Because heteroplasmy dictates phenotype, the reduction of heteroplasmy is potentially therapeutic. We identified pathogenic variants that increased the potential for formation of non-canonical G-quadruplexes (GQ) within mtDNA. The Leigh Syndrome (LS)-associated mt.10191T>C variant has a high probability of local GQ formation that was enhanced by the variant. Structural studies of mt.10191C-containing oligonucleotides confirmed the formation of GQ, and its interaction with the small molecule GQ-binding agent berberine increased GQ stability. The GQ formed at mt.10191 impeded polymerase processivity, and inhibition was enhanced by the mt.10191C variant. We applied a cyclical treatment of two GQ binding compounds, berberine or RHPS4, to primary fibroblasts from LS patients with heteroplasmic mt.10191T>C mutation. This treatment induced alternating mtDNA depletion and repopulation and was effective in shifting heteroplasmy towards the nonpathogenic allele, leading to an increase in complex I protein levels. This study demonstrates the potential for using small-molecule GQ-binding agents to induce beneficial shifts in mitochondrial heteroplasmy.

## Introduction

Mitochondrial diseases pose formidable challenges in diagnosis and treatment (1). The diseases are bigenomic, caused by pathogenic variants in either the maternally inherited mitochondrial genome (mtDNA) or the nuclear genome (nDNA) that affect mitochondrial respiratory complex function. Although nuclear-encoded defects predominate in infancy, most later-onset forms of mitochondrial disease are due to pathogenic variants within the mtDNA (2). Patients with disease due to heteroplasmic mtDNA mutations often have poor prognosis and we lack medications that offer specific utility in the treatment of mitochondrial disease (3–5).

Despite this, the development of genomic therapies for mitochondrial disorders due to pathogenic variants in mtDNA has a distinct theoretical advantage when compared to the modification of nuclear DNA. With few exceptions, patients are heteroplasmic, retaining some proportion of non-pathogenic mtDNA. The relative levels of wild type and variant-bearing mtDNA affect the timing, severity and range of symptoms encountered (6, 7).

The “shifting” of heteroplasmy towards an improved genotype, either by enhancing the replication of mtDNA lacking the pathogenic variant or by inhibiting the replication of mutation-bearing mtDNA, may be the key to the development of new therapies (8). Several studies have approached this problem through the nuclease-dependent cleavage of mtDNA containing pathogenic variants, a strategy that encourages the repopulation of cells by the residual, wild type mtDNA (9–14). Specific cleavage of the pathogenic-variant mtDNA effectively reduces the heteroplasmic load in cells derived from patients with disease or in animal model systems. However, despite the considerable promise of this strategy, heteroplasmy shifting through mtDNA cleavage has not advanced to clinical use. Hurdles to therapeutic development include the complexity and risk of widespread introduction of sequence-specific nucleases, the challenge of reaching all affected organs and tissues and the high number of target mitochondrial genomes per cell.

In this study we have developed a means of heteroplasmy shifting based on the impact of specific pathogenic mtDNA variants on the formation of G-quadruplex (GQ) structures within the mtDNA itself. GQs are a non-canonical secondary structure that forms in single-stranded DNA through Hoogsteen pairing between four homopolymeric guanine repeats that form a tetrad (15). The study of GQ structures has greatly expanded with the recognition that they have regulatory function at promoters and telomeric sequence (16).

Due to its asymmetric base composition, there is a natural tendency of the purine-rich heavy strand of mtDNA to form GQs. GQ formation within the mtDNA has been inferred from studies of mtDNA deletion breakpoints, which are enriched at regions with high GQ-formation potential (17). The existence of mitochondrial GQs has been directly inferred by the co-localization of GQ-binding compounds with mitochondria-specific markers (18). Attempts to characterize the full set of GQ-forming regions in human DNA identified regions of the mitochondrial genome by immunoprecipitation with GQ-specific antibodies (19). Another genome-wide approach based on differential massively parallel sequencing, known as G4Seq, also characterized regions of mtDNA that formed GQs (20). GQs are also posited to play a physiological role in balancing the expression of mitochondrial genes and mtDNA replication. Specifically, an RNA-based GQ regulates the transition from transcription to DNA replication *in vitro* by interfering with RNA polymerase (POLRMT) processivity, promoting a switch from mitochondrial transcription to mtDNA replication (21, 22). Taken together, these findings confirm the presence and relevance of GQs for mitochondrial gene regulation.

In this study, we identified pathogenic variants in mtDNA that enhance the predicted formation of GQs. Differences in GQ formation between mtDNA bearing the wild type and pathogenic variant alleles could be exploited to induce a heteroplasmy shift. Increased GQ formation at the pathogenic allele could act as a selective obstacle to DNA replication since stabilized GQ structures can block polymerase processivity (16). Such an approach would lead to heteroplasmy shift through the preferential replication of wild type mtDNA. In this study, we have explored the effect of a specific pathogenic mtDNA variant *in vitro* and in patient cell lines and evaluated the possibility of enhancing GQ-formation through the use of small molecules that bind and stabilize GQ structures.

## Results

### Evaluation of mitochondrial pathogenic variants and GQ formation

The GQ-forming potential (GQFP) of the entire mitochondrial sequence has been previously studied (23). We re-evaluated the sequence with a second predictive software to identify all potential GQ-forming regions (24). We then compared these regions to all variants annotated in the MITOMAP database that have a confirmed pathogenic designation (25). Pathogenic variants that overlapped with GQ-forming regions were considered further. We extracted all variants that altered the length of poly-guanine tracts on either the heavy or the light strand of mtDNA, because these mutations would potentially alter the ability of the mtDNA to form GQs.

We selected the variants that strengthened existing GQ predictions, where the pathogenic variant potentially allowed the formation of a more stable GQ than the wild type sequence (**Table 1**). We also identified all pathogenic variants that created overlapping GQ structures in the local sequence without increasing the maximal predicted score, since these could provide an alternative means to stabilize GQs. As was anticipated based upon the purine bias of the heavy strand of mtDNA, all of the identified pathogenic variants are T>C transitions on the light strand. The actual impact on GQ formation was through the complementary A>G transition, which lengthened homopolymeric guanine stretches on the complementary (heavy) strand.

**Table 1.**
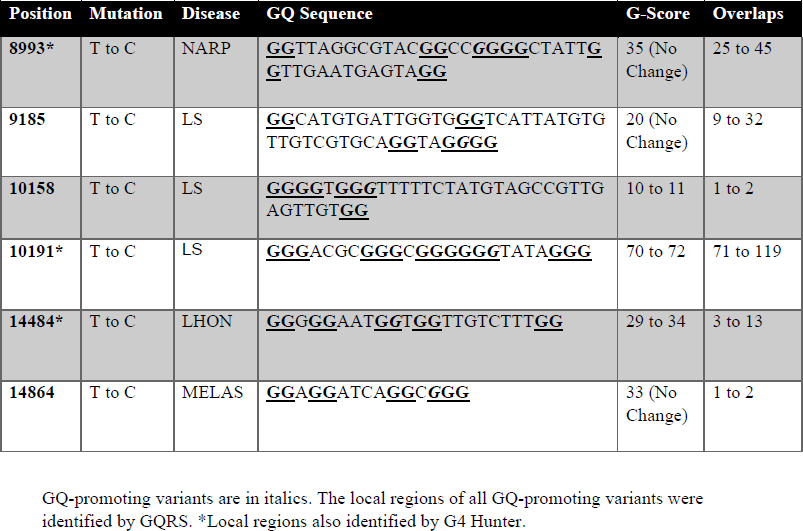

The mt.10191T>C pathogenic variant was selected for further study. Although exact incidence figures are not available for this mutation, it is viewed as a relatively common cause of complex I-deficient Leigh syndrome (LS) (26). All known patients with this mutation are heteroplasmic (26–28). The heavy-strand region proximal to mt.10191 has a high potential for GQ formation and the mt.10191C variant was predicted to increase both the number of potential GQs and their stability by extending a stretch of five guanines that is present in wild type mtDNA.

### Confirmation of GQ formation by mt.10191

We used circular dichroism (CD) and ultraviolet melting curve (UV-melting) studies to characterize GQ formation in the region proximal to mt.10191 *in vitro* and to determine the effect of the mt.10191T>C pathogenic variant (**Figure 1A**). The CD spectra for the mt.10191T and mt.10191C oligonucleotides had a trough at 240 nm and a peak at 262 nm, consistent with parallel GQ formation (29) (**Figure 1B)**. To determine the relative stability of the GQs formed by mt.10191T and mt.10191C, thermal denaturation profiles were obtained for the CD spectra by evaluation of the 262 nm maxima (**Figure 1C**). There was a rightward shift of the mt.10191C oligonucleotide thermal denaturation profile relative to mt.10191T, consistent with increased stability due to the pathogenic variant. We confirmed the relative stability of the GQs using UV-melting after folding of the oligonucleotides (**Figure 2**). In these studies, the melting temperature of mt.10191T was 65°C while the Tm for mt.10191C was 70°C, confirming the increased stability related to mt.10191C.

**Figure 1.**
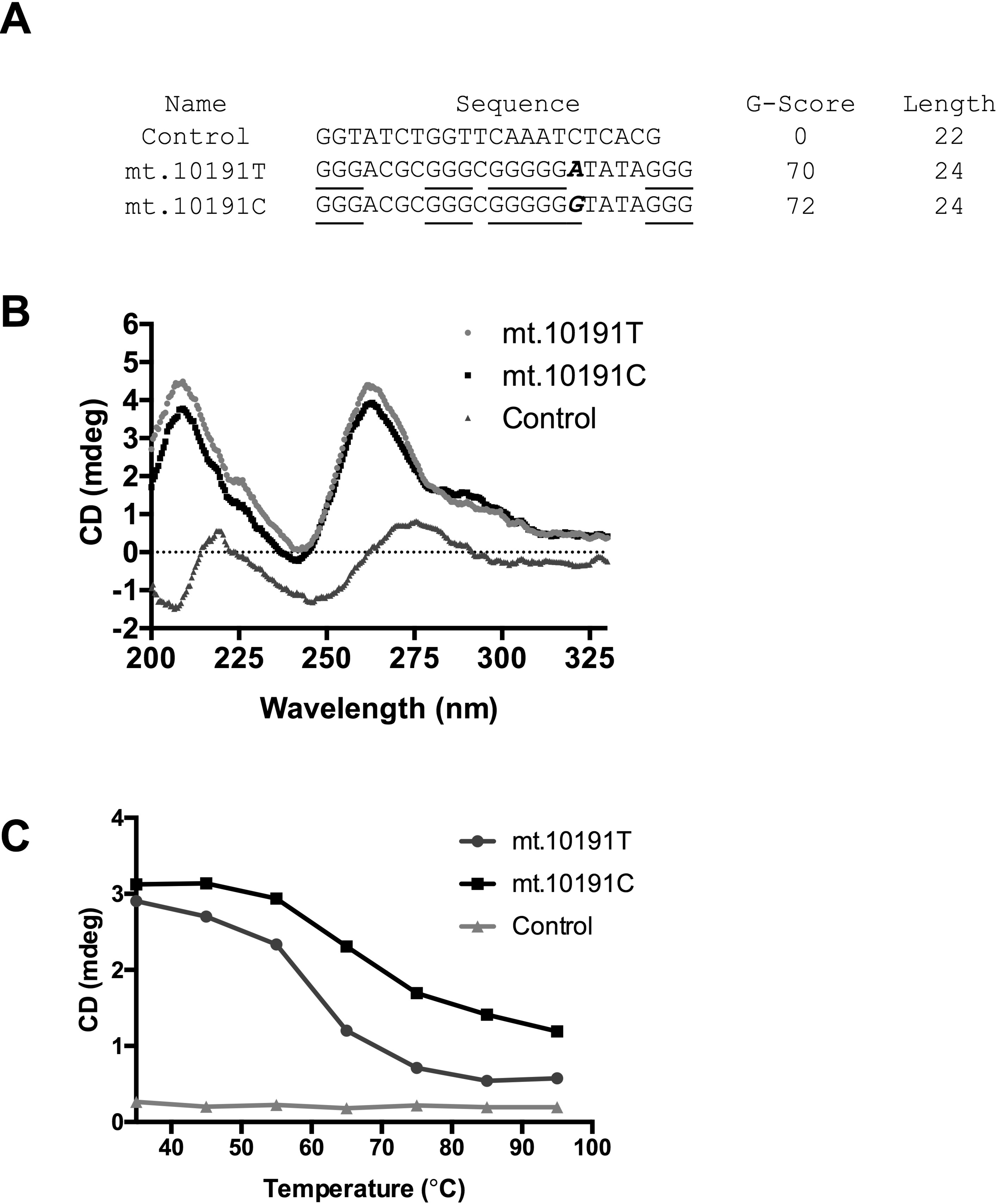
(A) Oligonucleotides used for CD and UV-M studies. Poly-guanine repeats longer than two are underlined. The location of the polymorphism is shown in bold italics in the context of the heavy strand region mt.10184-10207. G-Score is from QGRS (24). (B) CD plots after cooling under GQ-promoting conditions. The presence of the maxima at 262 nm and minima at 240 nm is characteristic of parallel GQ structures. (C) The thermal denaturation of the oligonucleotides was monitored at 262 nm.

**Figure 2.**
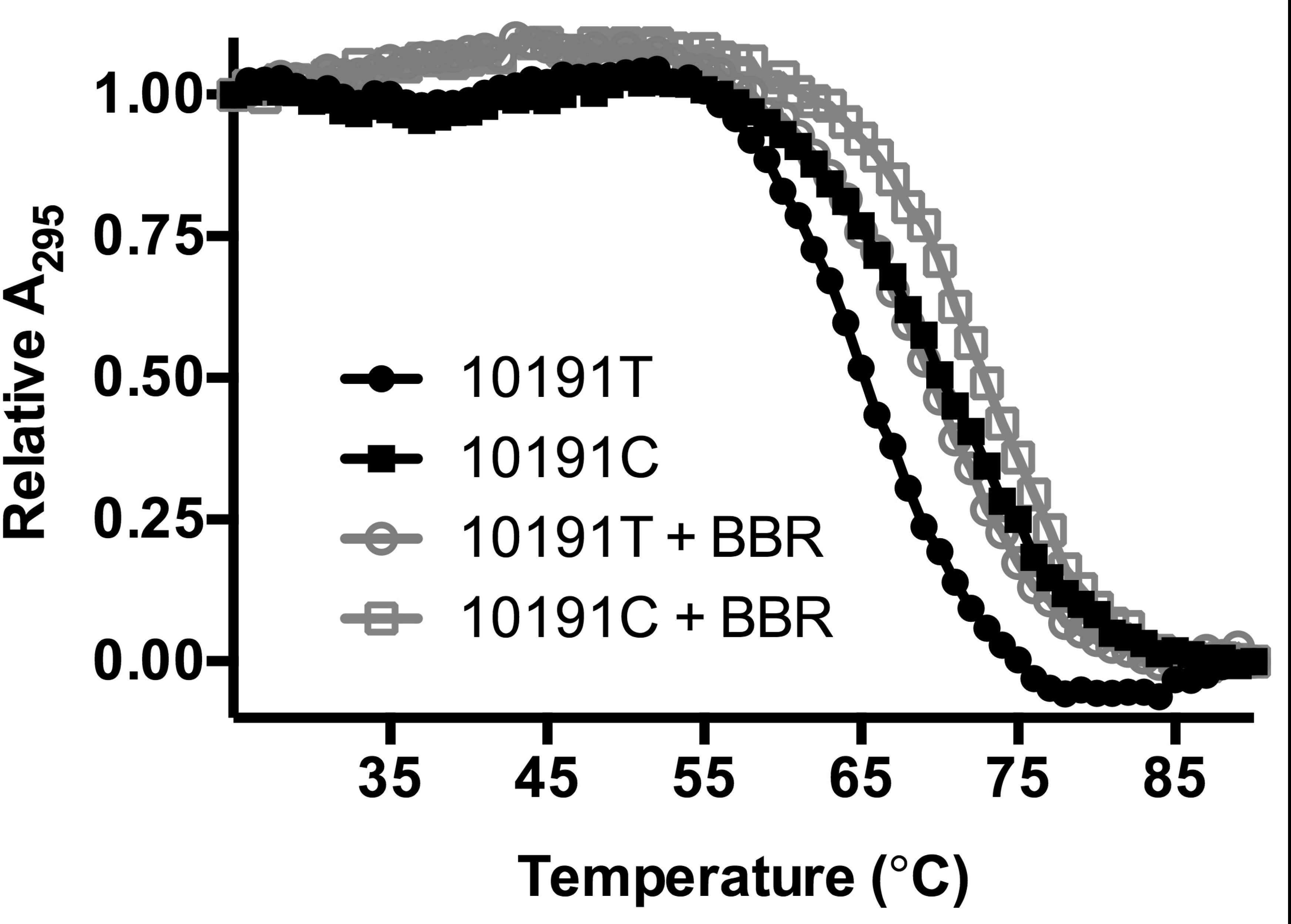
UV-Melting curves for mt.10191C and mt.10191T oligonucleotides (sequences as in Figure 1) in the presence or absence of 10 µM BBR.

We next tested whether the GQ structures formed by mt.10191C and mt.10191T were affected by the presence of GQ-binding agents (GQBA). GQBAs act by intercalating between the parallel tetrads of guanines formed by the GQ structure (16). We used the naturally occurring GQBA berberine hydrochloride (BBR), a plant alkaloid that is known to interact with telomeric GQ structures (30) but also localizes to the mitochondrial matrix (31). The addition of 10 µM BBR to previously folded oligonucleotides increased the Tm of both mt.10191T and mt.10191C by 4°C (**Figure 2)**. This confirmed that the region proximal to mt.10191 forms a GQ and interacts with GQBAs to achieve greater stability.

### Polymerase inhibition by GQ structures at mt.10191

GQ-GQBA interactions should potentially impact heteroplasmy through their effect on processive polymerases. In order to be effective in heteroplasmy shift, the mt.10191C allele must have a differential impact on DNA replication. To model this possibility *in vitro,* we performed primer extension assays with a labeled primer that anneals to the template DNA at a position 5’ to the GQ-candidate region and extended it to test the capacity of the region to interfere with processivity. We used the Family A polymerase member *Taq* as a model of mitochondrial DNA polymerase (POLG) activity (**Figure 3A**).

**Figure 3.**
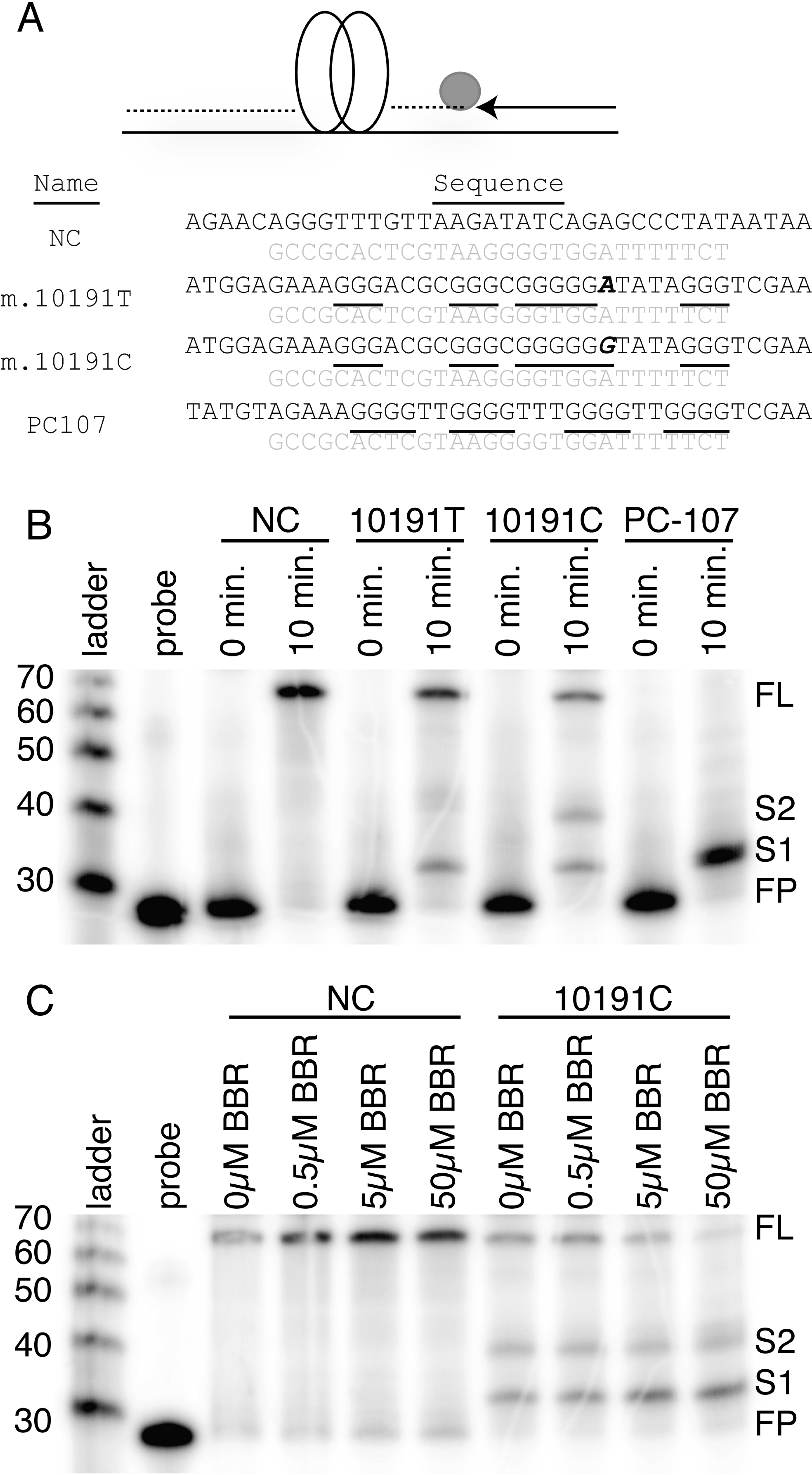
The capacity of *Taq* polymerase to extend through regions susceptible to GQ formation was tested by primer extension. The schematic of the experiment and oligonucleotides used are shown (A). The light grey nucleotides are complementary to the labeled primer, the poly-guanine tracts are underlined and the mutation is marked in bold. (B) Primers were extended over templates and resolved by denaturing gel electrophoresis. These included a negative control (NC) sequence that does not form GQ, the region complementary to mt.10150-10216 containing the wild type T allele (10191T), the same region with the C allele (10191C) and an artificial construct with optimized GQ formation (PC-107). 0 min marks the time before nucleotide addition. FL marks the expected size for the full-length runoff products (66nt.), FP marks the free probe and two stopped products S1 and S2 are indicated. The labeled products were isolated and separated with a single-strand DNA ladder. (C) The control and mt.10191C templates were extended in the presence of the labeled concentration of BBR.

First, we compared the extension through the mt.10191 region containing either the T or C allele to a non-GQ forming negative control template of the same size. A positive control sequence, which had been created to maximize GQ formation at the expected position of GQ-dependent inhibition of DNA synthesis (PC-107) (32), was also included. Primer extension templates were formed under GQ-promoting conditions (80 mM KCl) and we confirmed that these structures were formed by unimolecular interaction of poly-guanine tracts (**Supplemental Figure 1**). Extension of the negative control sequence proceeded to completion to form a runoff product when the labeled product was evaluated by denaturing electrophoresis (**Figure 3B**). As expected, the extension of the artificial PC-107 positive control was completely blocked at the 5’ end of the predicted GQ structure so that no full-length product was formed. The extension of oligonucleotides containing either the mt.10191T or mt.10191C alleles was marked both by the formation of GQ-truncated products and a partial loss of full-length extension. Interestingly, we repeatedly observed two clear termination products of different lengths in reactions on the mt.10191C template, suggesting the possibility of a second polymerase-inhibitory conformation for extension over the mt.10191C sequence.

Next, we evaluated whether extension over the GQ forming region could be affected by the presence of BBR. We compared the extension of the negative control sequence to that of mt.10191C. Progressive addition of BBR up to 50 µM did not impair *Taq* activity on the control template, but progressively inhibited processivity along mt.10191C until minimal full-length product was produced (**Figure 3C**). This demonstrated that the interaction of the GQBA BBR with the mt.10191 region impaired the replication of mtDNA in a manner responsive to the dose of GQBA.

### Alternative GQ structures are favored with the mt.10191C allele

The presence of a strong secondary inhibition on mt.10191C templates suggested that there was an alternative GQ conformation available when the template contained the pathogenic variant. The size difference in electrophoretic mobility between the S1 and S2 products (seen in **Figure 3**) suggested that the product formed at S2 was not due to a GQ that included the most proximal triplet of guanines, which is the first encountered by the polymerase in the assay (**Figure 4A**). Instead, the S2 product could form if the GQ began with the expanded block of guanines that included the pathogenic variant. Were this to occur, the six-guanine tract could become the two initial poly-guanine repeats of the GQ structure. This type of GQ structure is known as a zero-loop because there is no intervening sequence between two of the guanine tracts (33). This structure cannot be as readily formed by the mt.10191T allele because it lacks the expanded repeat of guanines. Instead, only two-stack or other alternative GQs can form which have reduced stability.

**Figure 4.**
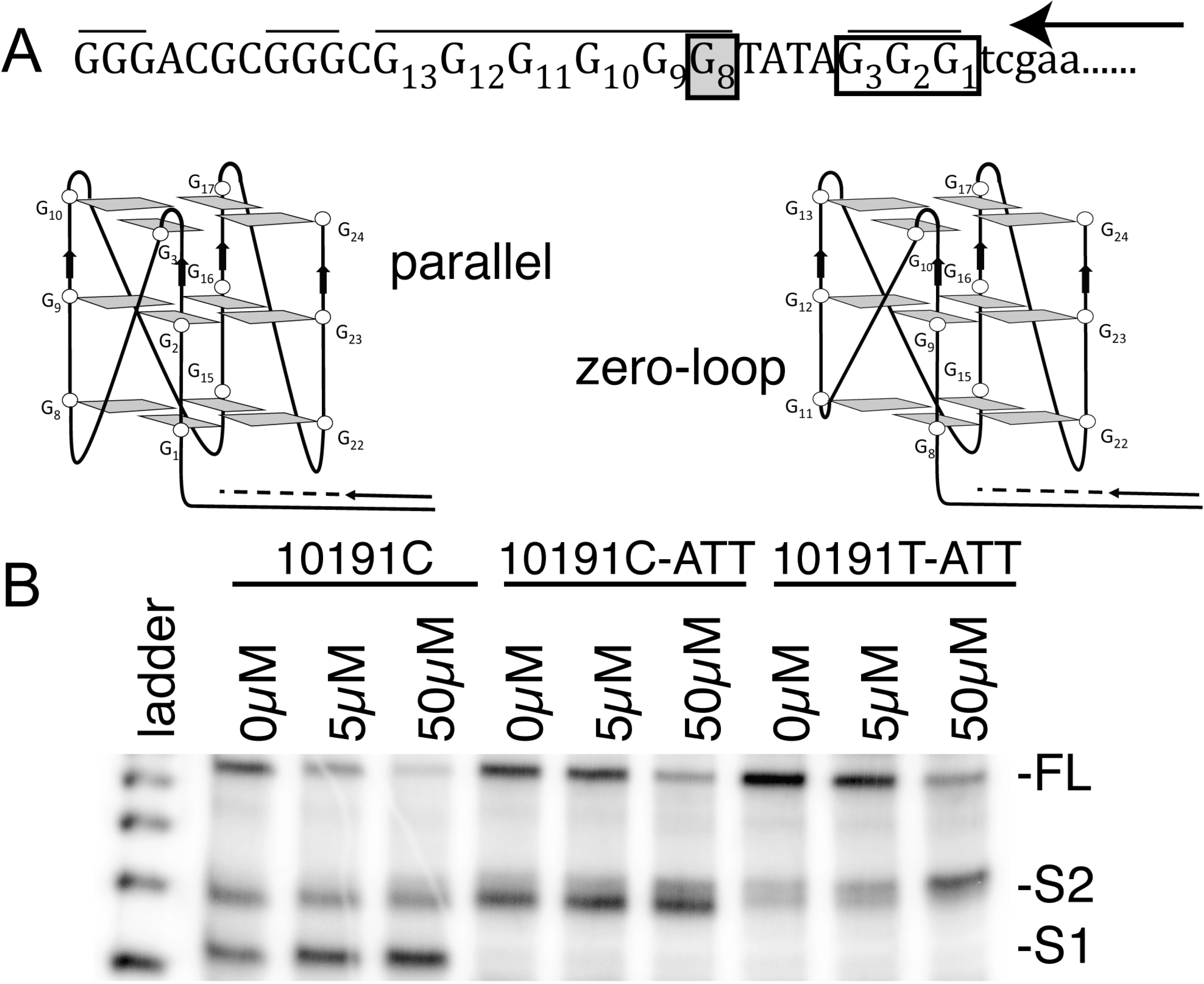
(A) Alternative GQ formation in the mt.10191 region. The mt.10191C mutation (grey box) extends the guanine repeat of block II. The clear box shows the three guanines mutated to create -ATT variants of the oligonucleotide. The mt.10191C allows the formation of a zero-loop, triple-stack GQ shown at right, which is unavailable to mt.10191T oligonucleotides because it lacks G_8_. The arrow indicates the direction of DNA replication. (B) Templates with or without a deletion of the first block of guanines (-ATT) were subjected to primer extension in the presence or absence of BBR.

To test this possibility, we designed sequences in which the most proximal guanine triplet (G_1_G_2_G_3_ of **Figure 4A**) was replaced by nucleotides that would not be included in a GQ structure (-ATT mutation). Studies of the UV-melting and CD properties of the mt.10191TATT and mt.10191C-ATT oligonucleotides confirmed that they are both capable of forming GQs (**Supplemental Figure 2)**. When comparing allele-specific effects, we observed that the melting point of mt.10191T-ATT is lower than mt.10191C-ATT, suggesting that the GQ formed by the pathogenic allele had greater stability.

The -ATT templates are unable to form the truncated product S1 and therefore allow specific testing of GQs that use the larger guanine tract twice. We evaluated this again using the primer extension assay. First, we confirmed that mt.10191C-ATT was unable to support the formation of the S1 product (**Figure 4B**). We observed that the 10191C-ATT template strongly supported the formation of S2. The addition of BBR to the extension assay increased termination and decreased the formation of full-length product in response to the dose. The size of the termination product formed by mt.10191C-ATT was identical to the S2 product formed by the natural template. This finding is consistent with the ready formation of a zero-loop, triple-stack GQ product by mt.10191C (33).

Primer extension in the presence of the -ATT mutation was also evaluated on the mt.10191T template. This template drove weaker termination at slightly greater length than the S2 product, and the response to BBR was again noted (**Figure 4B**). We speculate that the termination product of mt.10191T-ATT may be due to the formation of a two-stack GQ with reduced inhibition of polymerase activity, particularly in the presence of BBR, when the formation of the canonical GQ using four poly-guanine stretches is unavailable. We conclude that the formation of a zero-loop product at S2 enhances the capacity of mt.10191C to interfere with polymerase activity.

### Depletion of mtDNA by GQBAs

We hypothesized that GQBA must induce mtDNA depletion in order to result in heteroplasmy shift. The loss of mtDNA would provide a stimulus for mtDNA replication, which would be biased by the presence of stabilized GQ structures at the variant allele to produce more wild type genomes. We evaluated the feasibility of using GQBAs to inhibit the replication of mtDNA in primary human fibroblasts from healthy foreskin samples. We first confirmed that BBR localized to the mitochondria via fluorescence microscopy using a mitochondria-targeted dye (**Supplemental Figure 3A-C**). Next, dose response studies were performed to determine concentrations of BBR that depleted mtDNA without adversely impacting cell division. A concentration of 0.3 µM BBR depleted mtDNA by 35% without any loss of cells (**Supplemental Figure 4**) and was used for further experiments. The repopulation of mtDNA after depletion occurred rapidly with both BBR and intermittent treatment with GQBAs had no significant impact on cell division. This suggested that cycles of GQBA exposure could potentially be effective in heteroplasmy shift.

### Heteroplasmy shifting in patient cell lines

We tested the capacity of GQBA administration to induce heteroplasmy shifting using fibroblast cell lines derived from patients with a diagnosis of LS due to mt.10191T>C. Experimental replicates (n=4) from a patient with moderate (initially 43%) load of the mutation were treated cyclically for eight weeks by exposure to 0.3 μM BBR with three days of treatment alternating with four days in drug free media (**Figure 5**). The exposure to BBR induced an initial mtDNA depletion of 41% of the control level, which rebounded to 72% after removal of BBR from the media. Continued application of cycles of BBR forced waves of depletion to levels distinct from untreated cells followed by repopulation to that were not significantly different from control.

**Figure 5.**
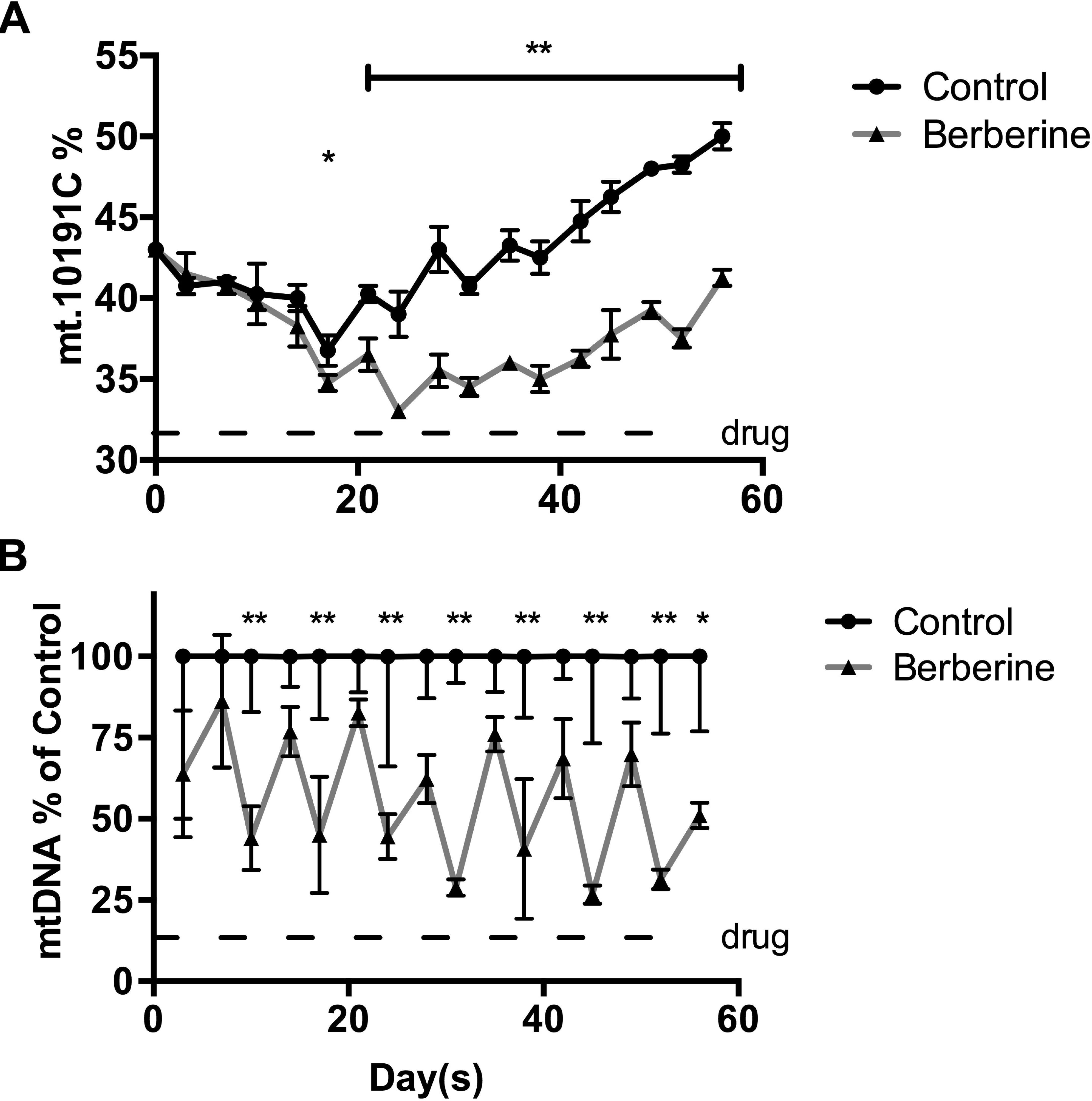
Fibroblasts derived from a patient with LS due to heteroplasmic mt.10191C mutation were grown in the presence of 0.3 µM BBR for 3 days, alternating with 4 days of BBR-free media, for 8 weeks, and are compared to the same cell line grown without BBR exposure (Control). A bar marks periods of drug administration. The proportion of mt.10191C (A) and the normalized ratio of mtDNA referenced to the Control (B) are shown. \**p*<0.05; \*\**p*<0.0001.

After two weeks of cyclic treatment, the heteroplasmy level of the pathogenic mt.10191C allele was significantly lower in cells treated with BBR compared to untreated cells. Over the course of the eight-week study, the difference in mt.10191C load in the BBR treated cells continued to increase as compared to untreated cells and the final difference in heteroplasmy between the treatments was 9%. The general upward drift of heteroplasmy in the untreated cells has been invariably observed with fibroblasts from this patient, and heteroplasmy drift is commonly observed in cell lines grown in culture (34).

One alternative possibility for heteroplasmy shift is that it is induced by waves of mitochondrial depletion and regrowth alone, with repopulation of the wild type allele through the selective advantage of having improved mitochondrial activity rather than through a GQ-dependent mechanism. To test this possibility, we repeated our studies with a paired exposure of cells to ethidium bromide (EtBr), an established technique for decreasing mitochondrial DNA copy number through intercalation into mtDNA. Notably, EtBr interacts weakly with GQ structures (35). Over a four-week exposure of heteroplasmic mt.10191T>C fibroblasts to EtBr and BBR, we induced equivalent depletion of mtDNA, but saw no corresponding change in genotype with EtBr (**Figure 6A&B**).

**Figure 6.**
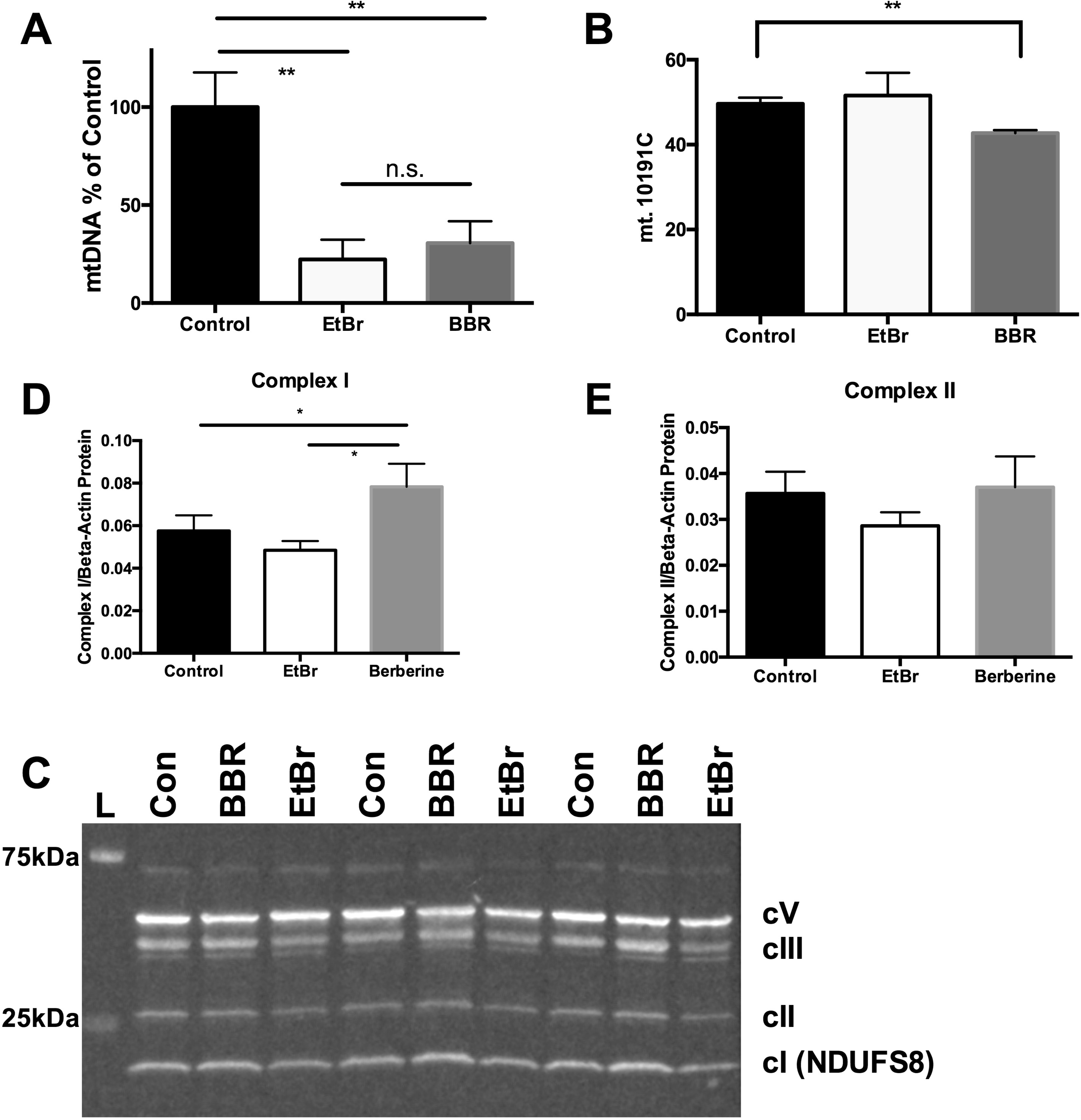
(A) mt.10191T>C fibroblasts were grown for 28 days with alternating cycles of 0.3 µM BBR or in the presence of 25 ng/mL EtBr. Mitochondrial depletion after drug exposure shows equivalent depletion in the EtBr and BBR conditions. (B) Heteroplasmy shift at 28 days shows a decline in pathogenic heteroplasmy in the BBR treated cell lines. (C) Western blotting with quantitation of replicates at 28 days of treatment. Differences are seen for complex I protein levels after BBR treatment (D) but not for complex II (E) or other complexes (quantitation not shown). \**p*<0.05; \*\**p*<0.0001.

We also evaluated whether treated patient fibroblasts had secondary indications of improved heteroplasmy. The mt.10191T>C mutation affects the ND3 protein, leading to decreased assembly of complex I of the electron transport chain (28). Following heteroplasmy shift, we observed higher levels of the complex I protein NDUFS8, but not components of other electron transport chain complexes, present in the BBR-treated fibroblasts when compared to either EtBr treatment or mock treatment (**Figure 6C-E**).

### Alternative GQBA RHPS4 induces heteroplasmy shift

Because BBR may have complex effects on mitochondrial physiology, it was necessary to confirm that the shift in heteroplasmy was specific to the GQ-binding activity of BBR. We evaluated the effect of a second GQBA on heteroplasmy shift. RHPS4 is a planar pentacyclic molecule that was developed because of its homology to GQBAs that inhibit telomerase (36). First, we demonstrated that RHPS4 localizes to the mitochondrial matrix as expected based upon its positive charge (**Supplemental Figure 3D-F**). Next, to confirm the potential of this compound to shift heteroplasmy, we evaluated RHPS4 using the polymerase extension assay along GQ-forming templates and demonstrated that it had a similar capacity to impeded polymerase along the region proximal to mt.10191 (**Figure 7A)**.

**Figure 7.**
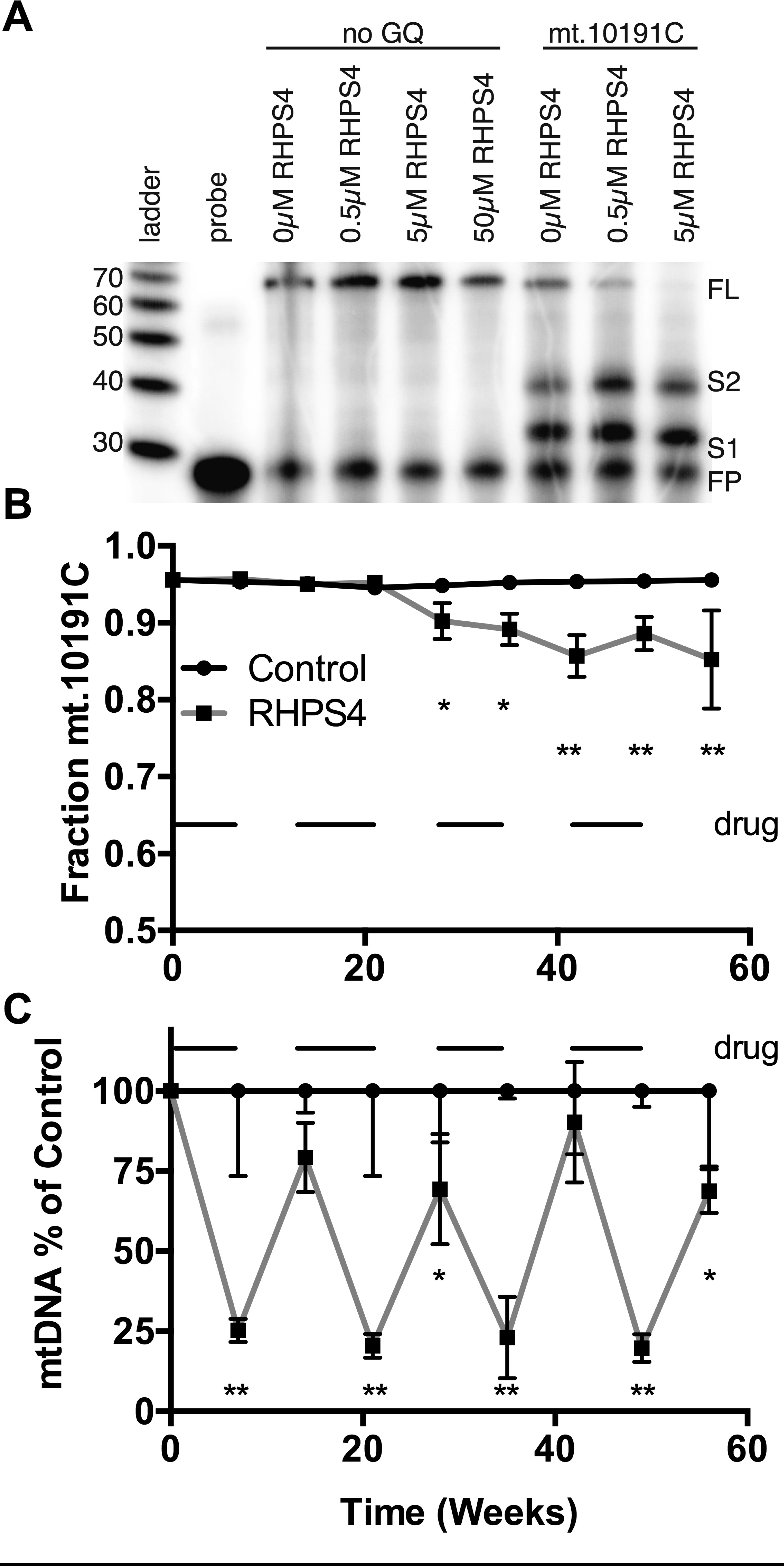
(A) Primer extension assays performed using the mt.10191C template responded with increased termination at S1 and S2 in the presence of increasing concentration of RHPS4. A template lacking GQ-formation potential was not affected. (B&C) Fibroblasts derived from a patient with high heteroplasmy mt.10191T>C mutation were grown in the presence or absence of 0.5 µM RHPS4 for 7 days, followed by 7 days of RHPS4-free media. A bar marks periods of drug administration. The proportion of mt.10191C (B) and normalized ratio of mtDNA (C) are shown. \**p*<0.05; \*\**p*<0.0001.

Using primary fibroblasts from a second LS patient with a higher initial degree of pathogenic mt.10191C heteroplasmy, we observed a decline in pathogenic heteroplasmy after treatment with 0.5 µM RHPS4 (**Figure 7B&C**). In the studies with RHPS4, we noted that shifts in genotype were most apparent during growth in the absence of the GQBA, and appeared after two cycles of treatment with RHPS4. We conclude that the treatment of mt.10191C-bearing patient cell lines with GQBAs has a favorable effect upon mitochondrial heteroplasmy.

### Studies of additional mutations

We evaluated heteroplasmy shift using patient-derived fibroblast cell lines bearing mutations that would not be susceptible to GQBA-mediated heteroplasmy shift, due to the lack of expansion of homoplasmic guanine repeats in the presence of mutation. Treatment of several cell lines that lacked GQ-formation capacity, such as mt.13513G>A, did not show differences between GQBA treated and untreated conditions (**Supplementary Table 1**), further suggesting that non-specific mtDNA depletion is not driving the observed effects.

## Discussion

Therapeutic heteroplasmy shift offers a means to reduce the symptoms of disease using the natural presence of the wild type allele in affected patients. Partial shifting of heteroplasmy in somatic tissues, without total elimination of the pathogenic mutation, is likely to be effective since it is well recognized that the load of pathogenic mutation correlates well with penetrance and severity and that heteroplasmy below a mutation-specific load may be incompletely penetrant for disease (6, 7). In this study, we explored whether an alteration in heteroplasmy could be induced through the binding of mtDNA by small molecules based upon the differential formation of GQ structures in pathogenic and wild type mitochondrial genomes. We focused on the pathogenic LS-associated mt.10191C variant, which is ideal for evaluating the potential for GQ-mediated heteroplasmy shift, because the pathogenic allele extends a poly-guanine tract on the heavy strand of mtDNA.

Our biochemical studies demonstrated that the pathogenic allele increases GQ stability in a region that already forms one of the most potent GQs within mtDNA. We observed that the GQs formed by this region are susceptible to pharmacological intervention using GQ-binding compounds *in vitro* and in patient-derived cells. The exposure of oligonucleotides containing this region to BBR increased its melting temperature and inhibited the processivity of DNA polymerase across the region. The polymerase inhibition was more pronounced for the pathogenic mt.10191C allele than for the wild type sequence, creating the possibility that treatment of heteroplasmic cells with GQBAs would bias the cell towards replication of the wild type allele. This effect of heteroplasmy shifting was demonstrated for two different GQBAs, with improvement of genotype after short duration, alternating cycles of treatment.

The observed heteroplasmy shift in mt.10191T>C cells may be principally driven through the formation of a zero-loop GQ structure. This structure is more stable in mtDNA bearing the mt.10191C allele because the length of the formed poly-guanine tract still allows the formation of a triple-stacked GQ. Although it has not been previously recognized that zero loop structures exist, recent studies of the hCEB1 minisatellite demonstrated that they are stable and form *in vivo* (33).

It is interesting that the zero-loop structure identified in hCEB1 was highly dependent upon an interaction with the GQBA Phen-DC3 (33). This may help to resolve an apparent paradox within our work. If GQs statically inhibit mitochondrial replication, the heteroplasmy of GQ-enhancing pathogenic mutations should always decline so that they are never observed. However, if in the case of mt.10191 differential replication of mtDNA is an emergent property in the presence of GQ binding agents, the pathogenic allele may be otherwise stable but treatable.

There has been a growing interest in the role of GQs in mitochondrial biology. An RNA GQ formed during the transcription of the heavy strand of the mitochondrial D-loop is crucial for switching between RNA synthesis and mtDNA replication (21, 22, 37). There is a high potential for GQ formation within mtDNA because of its biased nucleotide content, with considerable guanine richness on the heavy strand. In addition, the asymmetric mode of mtDNA replication, which remains to some extent controversial, involves intervals of single-stranding of the parental H-strand (38). TFAM, the ubiquitous mtDNA binding protein, has high affinity for GQ DNA (39). The Pif1 helicase, which is resident in the mitochondrial matrix, is stimulated by the presence of GQ and may interact and unwind the structures (40, 41). GQs may also bear responsibility for mtDNA deletions, as shown by the association of high-GQ potential regions with the ends of deletion breakpoints (17).

The manipulation of gene expression and DNA structure by GQs has been an area of medical interest because of the occurrence of GQ-forming structures in key promoters and in the human telomeric repeat (42). GQBAs, including RHPS4, have been developed for use as anti-cancer agents, based upon their potential to interfere with telomerase (36, 43, 44). It is convenient, however, that RHPS localizes to the mitochondrion. Studies of RHPS4 in animal models of cancer provide us with early data that the drug can be well tolerated in a whole animal model (44). Naturally occurring compounds with GQBA activity, including BBR, have been investigated as therapies for diverse conditions including diabetes, autoimmune disorders and cardiovascular disease (45). BBR is an activator of AMPK (46) and its overall effect on mitochondrial metabolism may be complex. It is proposed that BBR provides mitochondriogenic stimulus, and may increase mtDNA content in the face of hyperglycemia in a high-fat diet animal model (47). Some of these effects may be independently beneficial to patients with mitochondrial disorders, making BBR an appealing molecule for study. Other compounds may have superior activity against specific pathogenic alleles or may have superior side-effect profiles in clinical development. We are currently engaged in screening for molecules that may have improved characteristics for GQBA-mediated shift.

Heteroplasmy shift can be achieved by either promoting the replication of the wild type allele or by the selective inhibition of replication for the mutant allele. The data from our treatment of patient cell lines with GQBAs supports the latter mechanism, and are consistent with a model where GQBAs remain more tightly bound to the mutant allele after the removal of drug, converting the non-selective inhibition of mtDNA replication into a biased model. This approach is consistent with the most commonly reported approach to heteroplasmy shift, which relies upon sequence specific cleavage of mtDNA by mitochondrial-targeted nucleases.

The modification of heteroplasmy by small molecules may offer advantages in therapy development when compared to nuclease-dependent therapies. Many GQBAs are positively charged which will enhance their concentration within the mitochondrial matrix. The distribution of GQBAs to the targeted tissues such as brain, liver, muscle and kidney may also be more convenient than targeting multiple tissue types with nucleases. There may also be a reduced potential for off-target mutagenesis.

Further exploration of this mechanism is required before consideration of its potential in patient care. Our work focused on the region proximal to mt.10191, which is predicted to have extremely high basal GQ forming capacity. It will be informative to look at the additional mutations that impact GQ formation to see if there is a general utility to this strategy. Because GQ-mediated heteroplasmy shift relies upon the depletion of mtDNA, further study would be required to show that the depletion of mtDNA could be tolerated. Prior studies have suggested that extensive losses of mtDNA are required before the emergence of symptoms in patients with depletion syndrome (48, 49) and thus it is plausible that the strategy could be carried out safely. Studies in animal models can be used to evaluate the impact of mitochondrial depletion and the potential for off-target mutagenicity of GQBAs.

## Supplemental Data Legends

**Supplemental Figure 1** – Because GQs relevant to mtDNA replication are formed from the interaction of 4 guanine repeats on the same template strand, it is important to confirm that the primer extension experiments were conducted on unimolecular GQs, rather than on structures formed by the interaction of multiple copies of the longest poly-guanine tract of multiple templates. Primer extension reactions were formed by annealing labeled probes to the candidate template sequences under GQ-promoting conditions. The mobility of the primer-template hybrids was evaluated using native gel electrophoresis, without extension, with several variants of the template sequence. The **–ATT** variants replaced the most proximal guanine triplet to evaluate the potential for the large central poly-guanine repeat to act as two of the four sequences of the GQ. The **Tetra** template replaced all poly-guanines tracts except the large central polyguanine repeat, a structure that can only form intermolecular GQs between four strands.

(A) Native electrophoresis of templates without extension. Unimolecular GQ forms migrated faster than the control band and were observed when using the 10191T, 10191C, -ATT and positive control (PC-107) templates. The Tetra templates migrated slowly consistent with the formation of an intermolecular GQ created from four separate strands. As marked, the primer-template hybrids were isolated from the gel for extension.

(B&C) Primer extensions were performed on the indicated primer-template hybrids. The negative control sequence, which does not form a GQ, creates a full-length product. The rapidly migrating primer-template hybrids from T10191, C10191 and PC-107 form the S1 product, but not the full-length product. Notably, the rapidly migrating primer-template hybrids from mt.10191C–ATT templates form the S2 product (see overexposure panel D). The -Tetra forms also create a product at this size due to the formation of an intermolecular-GQ which blocks at the same point. A loading control made from a labeled oligonucleotide (LC) is shown. These results confirm that the mt.10191 oligonucleotides and their corresponding -ATT variants formed unimolecular GQs rather than intermolecular structures that are unlikely to form *in vivo*.

**Supplemental Figure 2** – The mt.10191C-ATT and mt.10191T-ATT oligonucleotides form GQs. (A) CD spectra of equimolar oligonucleotides with the mt.10191C oligonucleotide is provided as a positive reference. (B) UV-melting curves show that mt.10191C-ATT melting is unchanged (70 °C) when compared to the mt.10191C oligonucleotide. The mt.10191T-ATT oligonucleotide melts at 61°C, which is less stable than the unmodified mt.10191T (T_m_=65°C).

**Supplemental Figure 3** – Mitochondria in live primary human fibroblasts were stained with MitoTracker Deep Red (A & D). BBR, which auto-fluoresces green (B), was shown to colocalize with mitochondria: MCC M1 0.83 ± 0.18, where M1 = green channel (drug) and M2 = deep red channel (MitoTracker) (C). RHPS4, which auto-fluoresces both green and red (red channel shown pseudo-coloured green in E), was also shown to colocalize with mitochondria: MCC M1 0.87 ± 0.17, where M1 = red channel (drug) and M2 = deep red channel (MitoTracker) (F).

**Supplemental Figure 4** – Effect of increasing concentrations of BBR or RHPS4 on mtDNA depletion (A) and cell growth (B). RHPS4 exposure was for one week. mtDNA was quantified in reference to nuclear copy number and normalized to vehicle control. BBR exposure was limited to 3 days (C) because of poor growth after longer exposure at 0.3 µM (D).

## Materials and Methods

### Evaluation of mtDNA variants and GQ formation

We identified all GQ forming sequences (GQFS) in the revised Cambridge Reference Sequence (NC_012920) using QGRS (50). This was cross-referenced with all confirmed pathogenic variants in mtDNA identified in MITOMAP (25) to identify those that increased the number or maximum QGRS score of the tested region.

## Circular Dichroism and UV Melting Spectra

DNA oligonucleotides were purchased in purified form (Integrated DNA Technologies, Corelville, IA). Samples were prepared as 5 μM oligonucleotide in 10 mM K_2_HPO_4_ buffer with pH 7.5 and supplemented with 80 mM KCl (GQ buffer). To induce GQ formation, oligonucleotides were incubated at 95°C for 5 min and then allowed to cool gradually to room temperature overnight. Circular dichroism results were averaged over three scans from 200-330nm, 20nm/min scan rate, 8s response rate. CD melting curves used 10° increments from 25-95°C. Circular dichroism data was collected using a J-810 CD spectrometer (Jasco, Halifax, NS). For some CD spectra, 10 µM BBR was added following GQ formation and incubated at room temperature for 1hr before measuring CD spectra.

UV-melting spectra were measured at 295 nm over 25-90°C with 1°C/min rate and 2 min holding period using a Cary 300 UV-Visible Spectrophotometer (Agilent, Santa Clara, CA). All data were normalized to a buffer blank at 25°C.

### Primer Extension Assay

Primers were labeled with [**γ**-^32^P]-ATP (PerkinElmer, Waltham, MA) using 1 μL of T4 polynucleotide kinase 10 U/μL (Thermo Scientific, Waltham, MA) and purified using Micro Bio-spin columns (Bio-Rad, Hercules, CA). The primer was annealed to the extension template in GQ buffer, incubated at 95°C for 5 min and cooled gradually to room temperature overnight.

For the gel purification assay, samples were separated using an 8% non-denaturing gel supplemented with GQ buffer. The gel was exposed to X-ray film for 1 h to visualize band locations. Bands of interest were excised and DNA eluted in gel elution buffer (250 mM KCl, 10 mM Tris-HCl, pH=8.3 and 1 mM EDTA) overnight at 37C° with orbital shaking. Subsequently, DNA was precipitate with 100% ethanol at −20C° overnight and re-suspended in GQ buffer.

The extension reactions were prepared in a 20 μL reaction mixture containing GQ buffer, 2 μL of 10X *Taq* buffer, 0.2 μL of 5000 U/ml of *Taq* (New England Bio-Labs, Ipswich, MA), and 5 μL of template DNA/primer solution. Following the 0-minute sampling 0.4 μL of 10 mM dNTPs was added to the reaction mixture. For BBR titrations the primer and template were incubated with 0.5, 5, or 50 μM BBR at 37°C for 1hr prior to the addition of dNTPs, 10X *Taq* buffer and enzyme. Reactions were terminated using 20 μL of alkaline loading buffer (80% formamide, 10 mM NaOH, and 0.005% w/v bromophenol blue). Products were resolved on a 12% denaturing polyacrylamide gel. The dried gel was developed using a Typhoon Phosphorimager (GE, Boston, MA).

### Cell Culture and Drug Treatment

Primary fibroblasts derived from donors were cultured in low glucose DMEM (Gibco, Thermo Fisher Scientific, Waltham, MA) supplemented with 10% FBS (Gibco), 1 mM sodium pyruvate (Gibco) and 50 µg/ml uridine (Acros Organics, Thermo Fisher Scientific). Cells were seeded in 6-well plates at a density of 10^5^ per well and exposed to 0.3 µM berberine hydrochloride (Sigma-Aldrich, St. Louis, MO) for 3 days or 0.5 µM RHPS4 for 7 days. Drug was then removed and cells were allowed to recover for 4 days (BBR) or 7 days (RHPS4). Drug-recovery cycles were carried out for a total of 8 weeks (8 cycles for BBR and 4 cycles for RHPS4). Cells were harvested following each drug and each recovery cycle for mtDNA and genotyping analysis. Re-seeding at 10^5^ cells per well took place every 7 days.

### Isolation of DNA, Quantification of mtDNA and Genotyping of 10191 Allele

Total DNA was extracted from cells via DNeasy Blood and Tissue Kit (Qiagen, Hilden, Germany). mtDNA was quantified by measuring relative amounts of the mitochondrial gene MT-COX1 and the nuclear gene COX4I1 by qPCR on a QuantStudio 5 Real-Time PCR System (Applied Biosystems, Thermo Fisher Scientific), as described previously (51). Genotyping of the 10191 allele was carried out using a 2-probe allele specific qPCR system (Life Technologies, Thermo Fisher Scientific).

### Mitochondrial Localization of GQBA

Primary human fibroblasts were seeded in a black, high-binding µClear 96-well plate (Greiner Bio-One International, Kremsmünster, Austria). Cells were exposed to 1 µM RHPS4 for 18 h or 25 µM BBR for 6 h. Drugs were removed and 150 nM MitoTracker Deep Red FM (Invitrogen, Molecular Probes, Thermo Fisher Scientific) was added in the absence of serum for 30 min in the dark. Images were acquired using a Quorum WaveFX-X1 spinning disc confocal system (Quorum Technologies, Inc., Guelph, ON, Canada) BBR was imaged with a 491 nm laser and 515/40 emission filter, RHPS4 was imaged with a 561 nm laser and 594/40 emission filter, and Mitotracker Deep Red was imaged with a 642 nm laser and 670/40 emission filter.

Image analysis was performed using the Colocalization module in Volocity software version 6.3. Ten cells in 8-10 fields of view were analyzed per drug.

### Statistical Analysis

Data were analyzed via two-way ANOVA with Sidak post-test. Statistical significance was set at p<0.05.

